# Overexpressing the IPT gene improves drought tolerance and nutritional value of tropical maize (*Zea mays* L.)

**DOI:** 10.1101/2023.08.14.512900

**Authors:** Rose Mweru Muruo, Shem B. Nchore, Richard O. Oduor, Mathew Piero Ngugi

## Abstract

Drought stress poses a significant threat to crop productivity, making the development of drought-tolerant crops a priority. The impact of drought on grain yield loss varies significantly, ranging from 10% to 76%, depending on the specific stage of occurrence and the severity of the drought. In this study, we investigated the effects of introducing the pSARK::IPT transgene on the drought tolerance and nutritional composition of successive generations of tropical maize. Towards this goal, we screened different generations of maize plants by genotyping PCR, exposed them to long term drought stress and analysed several drought stress markers and nutritional profiles of the plants. Our results demonstrated that the pSARK::IPT transgene was present in 4 successive generations of maize plants. Under drought conditions, transgenic maize exhibited higher relative water content, and delayed senescence compared to wild-type plants. Additionally, transgenic plants showed increased levels of total chlorophyll, chlorophyll a, and chlorophyll b, indicating improved photosynthetic activity under water deficit. Our study also showed that IPT-transgenic plants produced substantially higher yields and demonstrated enhanced nutritional value compared to wildtype plants when grown under well-watered conditions. Further research is warranted to investigate the underlying molecular mechanisms involved in these improvements and assess the performance of pSARK::IPT maize under field conditions.

## Introduction

Maize (*Zea mays* L.) is a type of grain that belongs to the Poaceae family (1). Globally, maize has been estimated to occupy about 168 million hectares, a production of 854 million tones and productivity of about 5,000 kilograms per hectare (2). In Sub-Saharan Africa (SSA), its dependency as a source of food income and livelihood is over 80% (3–6). In Kenya, maize represents at least 51 % of all the staple food crops grown. The consumption rate in Kenya is approximately 103 kg/person/year. Kenya is regarded as a great maize consumer with production shortages (7).

Water is the most limiting resource of agricultural production (8). Maize is third in high water requirement after sugarcane and rice. Frequent water deficit resulting from sporadic rainfall as well as soils with low water holding capacity causes a substantial drop in maize yield throughout the tropics (9). It is anticipated that climate change will cause great impacts on water availability for crop production in the coming years. The United Nations predicted that the world population will rise by more than tenfold in 250 years (10). Rapid population growth obstructs efforts to increase income, guard livelihoods and lessen food shortages, predominantly in rural areas where food insecurity is often most startling (11). More than 80% of Kenyan land is considered arid and semi-arid (12) thus making the country food insecure.

Drought compromises stomata function impairs gas exchange and which leads to overproduction of reactive oxygen species (ROS) and the development of oxidative stress (13). Moreover, water deficit inhibits cell division, expansion of leaf surface, growth of the stem, and proliferation of root cells (14). Drought tolerance can be referred to asis the ability of a plant to exhibit the lowest yield loss in a water-deficient environment comparative to the highest yield in an environment with adequate water for crop growth (15).

Different maize varieties that are drought tolerant have been developed via recombinant DNA technology with a focus on various genes that target various biochemical properties in plants. There are various genes involved in drought tolerance that have been extensively studied including *i*I*sopentyl transferase*. This gene that encodes *Isopentyl transferase* enzyme which catalyzes a key step in the biosynthesis of Cytokinin (CK) (16,17). This is an important plant hormone that modulate growth and development in plants as well as regulate plant response to water-deficit stress. It has been reported that CK signaling acts as an intercellular communication network that helps mediate plant stress response during drought stress by interacting with other types of phytohormones and their regulating pathways (18).

Leaves provide a photosynthetic environment for fixation of carbon dioxide for during carbohydrate synthesis. Senescence is closely linked to variations in expression profiles of a number of genes with concomitant deterioration of cells and nutrient recycling (19). As the plant organs senesce, biomolecules are degraded and the nutrients recycled to actively growing regions (20). Leaf senescence and abscission during dehydration stress enable plants to reduce the size of their canopy. Perennial plants use this mechanism for their survival as it enables them to finish their life cycle in water-limiting environments. Nonetheless, this decreases the yields causing financial loss to farmers (21,22).

Successful genetic engineering necessitates that transgenes exhibit stable expression which must be passed on to subsequent generations in predictable pattern (23). Transgene inheritance and expression levels are highly dependent on various factors, such as transformation methodologies, promoter strength, insertion sites and copy numbers of the transgene (24). Analysis of transgene expression has been conducted on various staple crops like barley (*Hordeum vulgare* L.) (25), maize (*Zea mays* L.) (26), wheat (*Triticum aestivum* L.) (27) and rice (*Oryza sativa* L.) (28). In the majority of the studies, silencing or loss of the transgene was reported.

Many studies have been done and more are still ongoing on the development of transgenic maize that confers tolerance to drought stress. *Isopentenyl transferase* gene driven by different promoters has been used to transform maize (29,30). This study therefore focused on transgenic maize generated at the plant transformation laboratory of Kenyatta University expressing *IPT* transgene driven by senescence-associated receptor kinase promoter (SARK). The study sought to ascertain whether transgenes are passed over to subsequent generations, determine whether the transgenic maize shows tolerance to drought as well as determine differences or lack thereof in nutritional profiles of transgenic and non-transgenic maize. The findings of this study proposed that the transgenic germplasms should advance to field trials for assessment as a long-term solution to food security in maize-dependent countries.

## Materials and Methods

### Plant material and growth conditions

Maize variety CML 144, transformed with IPT gene under the control of senescence-associated receptor kinase promoter (pSARK) (31)was used. Five seeds per pot replicated 3 times were potted in soil mixed with manure at a ratio of 2:2. The experiment was set up in a completely randomized design. The plants were maintained in the glasshouse at a temperature of 40/26 ±2 °C (day/night), relative humidity of 60±5 % and 16 h /8 h (light/dark) photoperiod. Untransformed CML 144 was used as the wild-type control throughout the experiment.

### DNA extraction and PCR analysis

Genomic DNA was extracted from three-weeks old leaves according to the CTAB method (32). To confirm the presence of the transgene, PCR analysis was carried out using gene-specific primers for the *IPT* gene and *PMI* gene, the selectable marker gene. The PCR amplification was conducted as per the previously established procedures (33). My Taq PCR master mix (Bioline) that included buffer, magnesium chloride and deoxy-nucleotide triphosphate primers and 1µl DNA template adjusted to a total volume of 25µl. The PCR machine (Eppendorf AG 22331, Hamburg, Germany) was programmed as follows; initial denaturation temperature of 95 °C for 5 min followed by 35 cycles each of 95 °C for 15 seconds, annealing at 55 °C (IPT) and 53 °C (PMI) for 30 seconds, initial elongation at 72 °C for 1minute, final extension at 72 °C for 7 min and held at 4 °C to infinite. The amplicons were resolved on 1.5 % (w/v) TAE agarose gel and run at 100 V for 45 minutes. PCR-positive plants were selected for drought stress assays. The oligonucleotide sequences used in PCR are shown below:

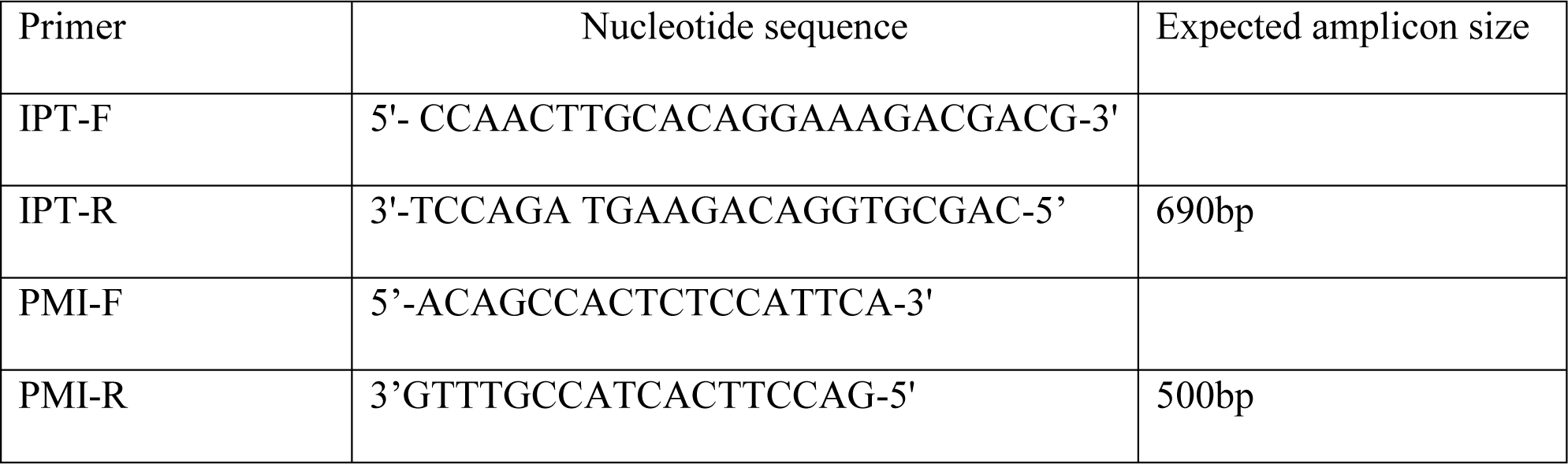

### Drought stress assays

The experiment was arranged in a completely randomized design in the glasshouse. The optimum amount of water for each plant was determined according to the method (34). The plants were watered daily with 2 L of tap water and maintained in the glasshouse. Sixty days post sowing, a cycle of drought conditions was induced by withholding water for 21 days. A control set was maintained by continuously watering the plants daily. After 21 days of water deficit, the plants were watered daily up to maturity. Sampling was done at days zero, seven, fourteen and twenty-one of the stress periods as well as three days after rewatering.

### Drought stress markers assays

#### Leaf relative water content (RWC)

To determine RWC, sampling was done at days zero, seven, fourteen and twenty-one of the stress periods as well as three days after rewatering. Three samples of leaves (3cm × 4cm) were excised from each experimental plant. The leaf relative water content was determined according to previously described method (35). The relative water content (RWC) was calculated according to the formula by (36).

#### Chlorophyll content

Chlorophyll was extracted from leaf 5 of each experimental plant using the dimethyl sulfoxide (DMSO) method (37). The optical density (OD) of the extract was measured at 649 and 665nm (722N Visible Spectrophotometer, EVERICH MEDCARE LTD, Nanjing, China), calibrating to zero with pure DMSO. Measurements and calculations were performed according to previously described methods (38).

#### Leaf senescence

Leaves showing at least 10% yellowing of the leaf blade were considered senesced. The number of senesced leaves per plant was counted at the 7^th^, 14^th^ and 21^st^ of the drought period. Additionally, a detached leaf assay was conducted as per previously described methods (52). Photographs were taken at three-and seven days post-incubation.

#### Antioxidant enzyme activity

Fresh leaf tissues were collected from leaf 5 of stressed and well-watered plants of both pSARK::IPT transformed and wild type plants. The crude leaf extracts were prepared according to a previously established method (39). The supernatants were refrigerated ready to be used to determine the various antioxidant enzyme activities.

Catalase activity was determined from leaves according to previously described methods(39). The decomposition of hydrogen peroxide was followed by a decrease in absorbance at 240 nm in a UV/Vis spectrophotometer (Specord 200 Analytik Jena). The extinction coefficient of hydrogen peroxide (40 mM^-1^ cm^-1^ at 240 nm) was used to calculate enzyme activity that was expressed in terms of milli-moles of hydrogen peroxide per minute per gram of fresh weight.

The Ascorbate peroxidase activity was assayed from maize leaves (40). The enzyme activity was determined from the decrease in absorbance (Specord 200 Analytik Jena) at 290 nm due to the oxidation of ascorbate in the reaction. The drop in absorbance was recorded for three minutes. The extinction coefficient of 2.8 mM^-1^ cm^-1^ for reduced ascorbate was used in calculating the enzyme activity which was expressed in terms of milli-mole of ascorbate per minute per gram fresh weight.

### Determination of selected agronomic parameters

#### Leaf rolling, plant height and leaf number per plant

The number of rolled-up leaves and the severity of rolling were scored per plant and recorded at 3 pm (the hottest part of the day) at day 0 and day 3 of stress. A leaf rolling scale with scores of 1-5 was developed and used to assess severity where 1 was considered normal (no rolling) while 5 was considered severely rolled up. Plant height was measured from the soil level to the tip of the uppermost fully expanded leaf at the end of the stress period. The total number of leaves was counted manually in each plant at the end of the stress period and the average was recorded as the number of leaves per plant.

#### Roots and shoots weight

At physiological maturity (120 days post sowing), plants were cut at the soil level. The shoots and shoots were weighed separately after removing the comb. The weight was recorded individually in grams per plant. Dry weight was determined by drying in the oven to a constant weight at 70 °C for 48 h. The cob was dried separately after removing the seeds.

#### Anthesis-silking interval (ASI), ear length, seed weight and seed number per plant

Days to anthesis were determined as the number of days taken by the plant from germination to the day the tassel started pollen shedding. Days to silking were determined as the number of days a plant had taken from germination to the day at least a single silk had emerged from the ear sheath. Anthesis-silking interval (ASI) was determined as the difference between the number of days to silking and the number of days to anthesis. The length of the ear was measured from the base to the tip and the value was recorded. The number of seeds per plant was counted manually and recorded as seed yield per plant. Additionally, the weight of 10 seeds was computed on a weighing balance and the value was recorded as well.

### Proximate analysis of mature maize kernels

Moisture content was previously described method (41) was used to estimate moisture content in the ground sample. Crude ash content was determined according to an already established method (42) Crude fat content was estimated by the soxhlet extraction method (43). Crude fibre content was determined by sequential acid and alkali hydrolysis method according to (44). To estimate crude protein content in the sample, the method described by (45).The nitrogen content was converted to proteins as shown below;

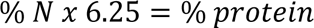

## Results

### Molecular analysis using PCR

To confirm the presence of IPT transgene in the plants, screening through PCR was conducted on T3 and T4 generations of *Zea mays* L. Fragments of 500bp and 690pb were amplified with PMI and IPT primers as shown in Fig 1. In T3 generation, three out of seven plants had the T-DNA while in T4, one out of seven plants had T-DNA. A total of 45 plants out of 125 events screened were positive for the T-DNA.

**Fig 1.**
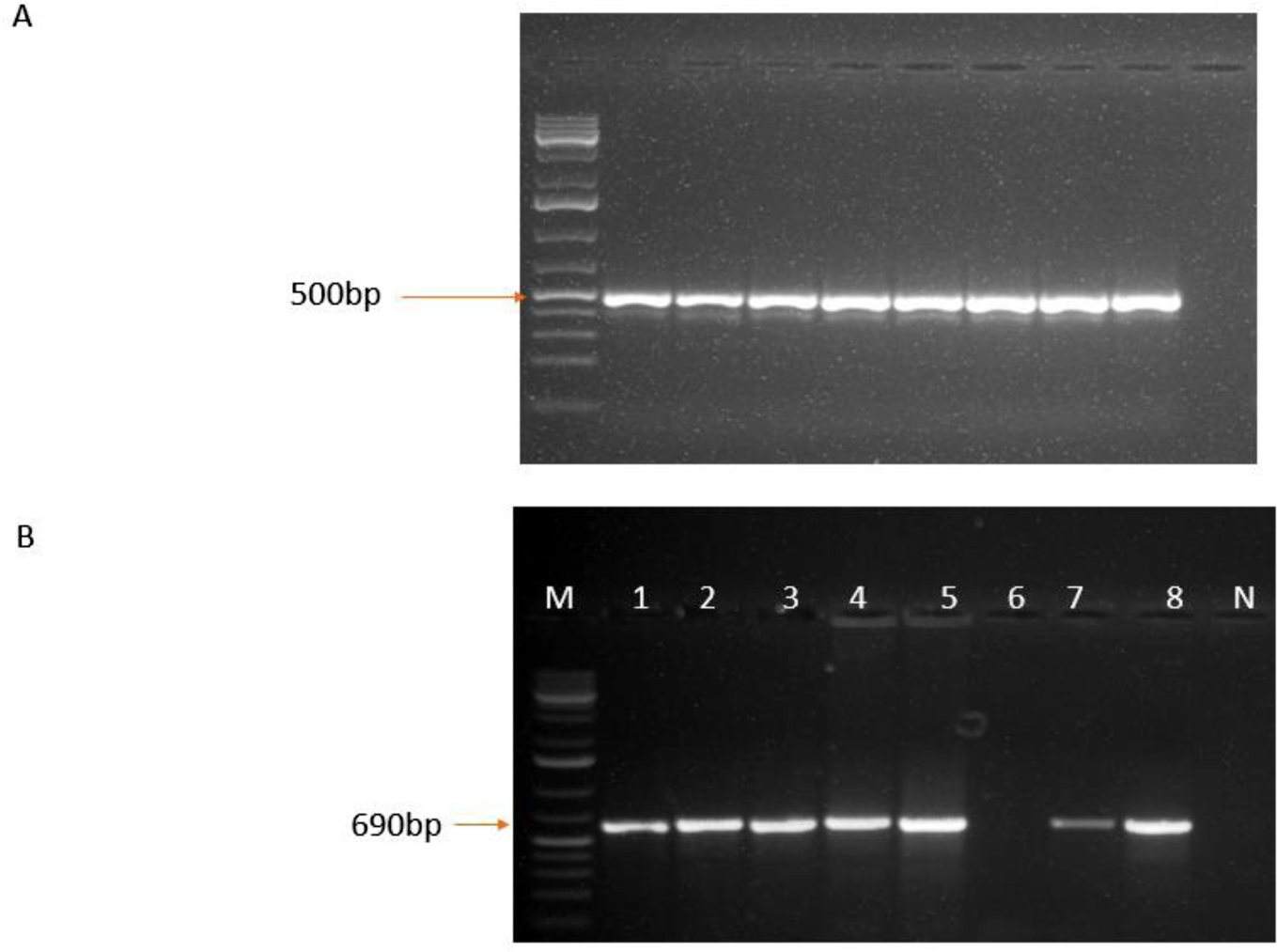
PCR products of transformed maize. **A**: PCR products (1–8) for detection of PMI gene and IPT gene **(B)** from T4 generation; M: 1kb plus DNA ladder (O’GeneRuler™ - ThermoFischer), N: No template control

### Effect of drought on leaf relative water content

Leaf relative water content (RWC), which was measured at different times points (0, 7, 14, 21 days) in leaf seven, revealed that there was a decrease in RWC in both wild type and pSARK::IPT maize plants upon exposure to drought stress. We found that the pSARK::IPT maize had significantly higher RWC than the wild type at all sampling points except at day zero (Fig 2). The RWC decreased from 79.10 % to 8.01% in wild-type maize after 21 days of stress whereas, in pSARK::IPT maize, the decrease was from 71.20% to 43.64%. Upon re-watering for 3 days, there was a 91% recovery in pSARK::IPT while the wild type had 10% of the RWC before dehydration stress (Fig 2).

**Fig 2:**
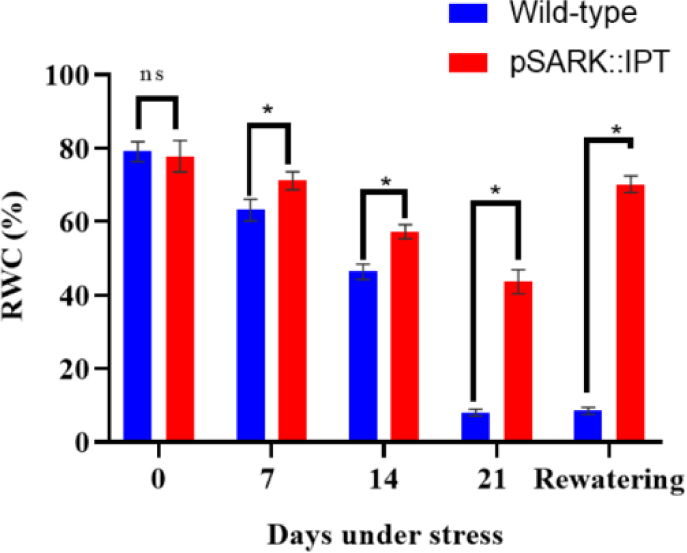
Leaf relative water content measured in 60-days old pSARK::IPT and wild type maize. Variation in RWC between pSARK::IPT plants and wild type. The results (n=15) are expressed as the mean ±SE. Graphs marked with asterisk (*) indicate that their means are statistically different from each other according to Student’s t-test at 95% confidence interval.

Additionally, leaf rolling was observed during the hottest part of the day (3 pm) before and during the stress period (Fig 3). Severe leaf rolling was observed in wild type maize within three days of dehydration stress, while in pSARK::IPT maize, leaf rolling was mild (Fig 3). The number of rolled-up leaves at day 0 and day 3 was statistically significant between the two groups (Fig 3). About 80 % and 12.22 % of leaves per plant were rolled up in wild type and pSARK::IPT plants respectively at day 0, while after 3 days of stress, rolled leaves were 83.82% in wild type and 27.6 % in pSARK::IPT maize. After seven days of stress, both groups exhibited leaf rolling.

**Fig 3.**
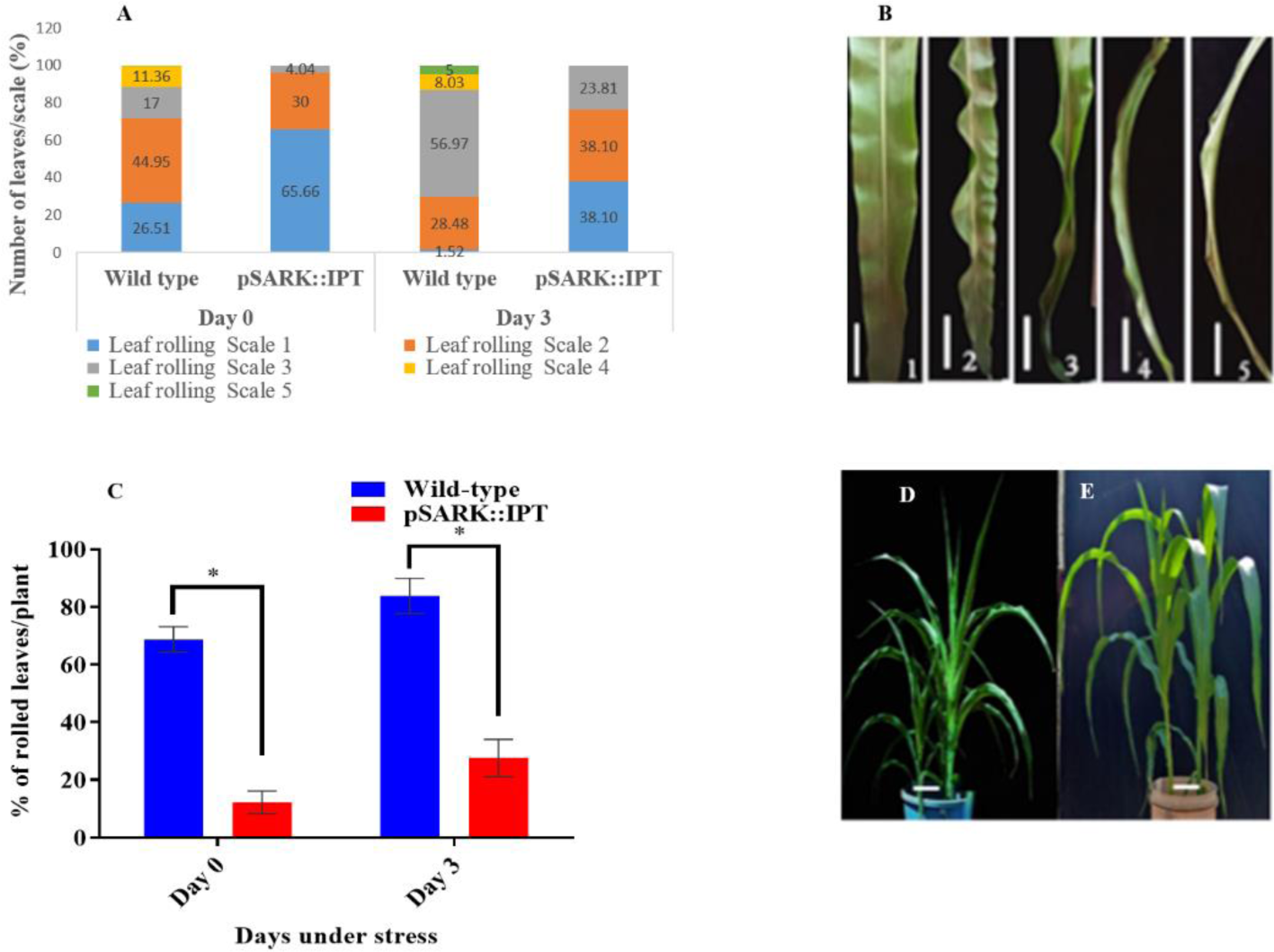
Leaf rolling in wild type and pSARK::IPT plants under drought stress. **A**: Severity score of leaf rolling in pSARK::IPT and wild-type plants under drought stress. **B**. Leaf rolling scale with 1 representing no rolling and 5 showing severe rolling. **C:** Number of rolled leaves per plant in wild type and pSARK::IPT plants at days 0 and 3 of drought stress. **D and E**: Leaf rolling in wild type and pSARK::IPT maize after 3 days of dehydration stress respectively. The results (n=15) are expressed as the mean ±SE. Bars marked with asterisk (*) indicate that their means are statistically different from each other according to Student’s t-test at 95% confidence interval. Scale bar = 1 cm.

### Effect of drought on leaf senescence

Leaf senescence, characterized by leaf yellowing, showed that the number of senesced leaves was significantly higher in wild type than in pSARK::IPT plants (Fig 4R). After 7 days of stress, 16.89 % of the leaves in pSARK::IPT plants were senesced in contrast to 72.92 % in wild type plants. By the 14^th^ day of stress, about 92.6 % of wild type leaves had senesced compared to 47.6 % in pSARK::IPT. At the end of the stress period, almost all leaves in wild type had senesced (98.4 %) unlike in pSARK::IPT plants where only 58.4% showed senescence (Fig 4R). Leaf 5 of all plants in both groups was closely monitored and the image analysis showed slower progress of senescence in pSARK::IPT plants compared to wild type under drought stress (Fig 4A-Q). Further, induction of senescence in the dark showed a delay in pSARK::IPT plants compared to wild type (Fig 4S-X).

**Fig 4.**
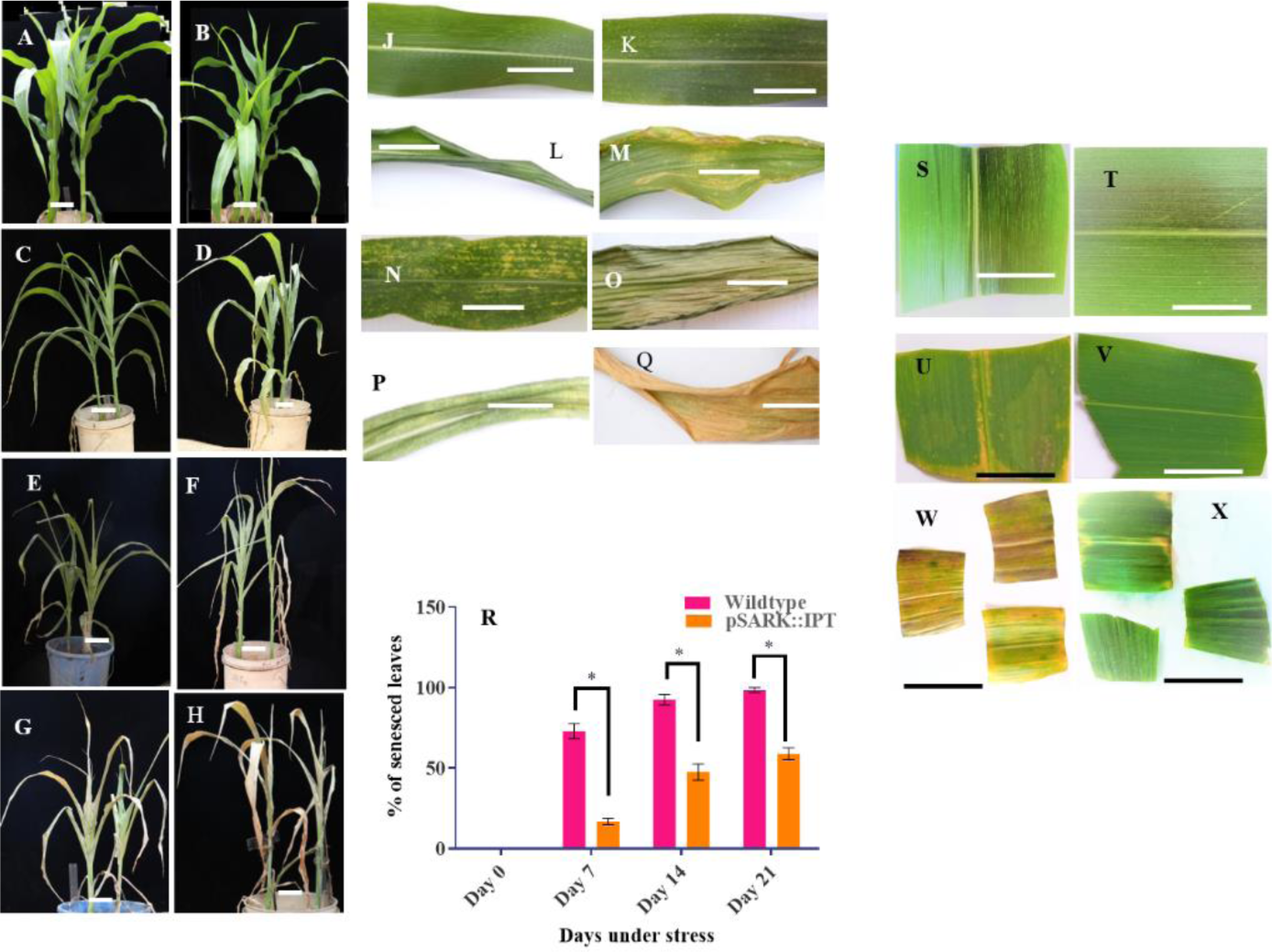
Progression of senescence in wild type and pSARK::IPT plants. **A, C, E and G**: Progression of senescence under drought stress in wild type plants at day 0, 7, 14 and 21 respectively. **B, D, F and H**: Progression of senescence in pSARK::IPT plants under drought stress at day 0,7,14 and 21 respectively. **J, L, N and P**: Progression of senescence in leaf 5 of wild type plants. **K, M, O and C** Progression of senescence in leaf 5 of pSARK::IPT plants. **S, U and W**: Induction of senescence in the dark in leaf 7 of wild type plants at days 0, 3 and 7 respectively. **T, V and X**: Induction of senescence in the dark in leaf 7 of pSARK::IPT plants at day 0, 3 and 7 respectively. The results (n=15) are expressed as the mean ±SE. Bars marked with asterisk (*) indicate that their means are statistically different from each other according to student’s t-test at 95% confidence interval. Scale bar = 1 cm.

### Effect of drought on chlorophyll a, b and a: b ratio

Leaf senescence is characterized by chlorophyll degradation. Chlorophyll a, the primary pigment for photosynthesis, was non-significantly higher in pSARK::IPT maize than in the wild type up to the 14^th^ day of the stress period (Fig 5). However, significant differences were observed between the two genotypes after 21 days of stress (2.23mg/gfw in pSARK::IPT and 1.73 mg/gfw in wild type). The pSARK::IPT maize and the wild type differed significantly after rewatering the plants for three days with pSARK::IPT plants having 1.73 mg/gfw chlorophyll while wild type had 0.45 mg/gfw of chlorophyll.

**Fig 5.**
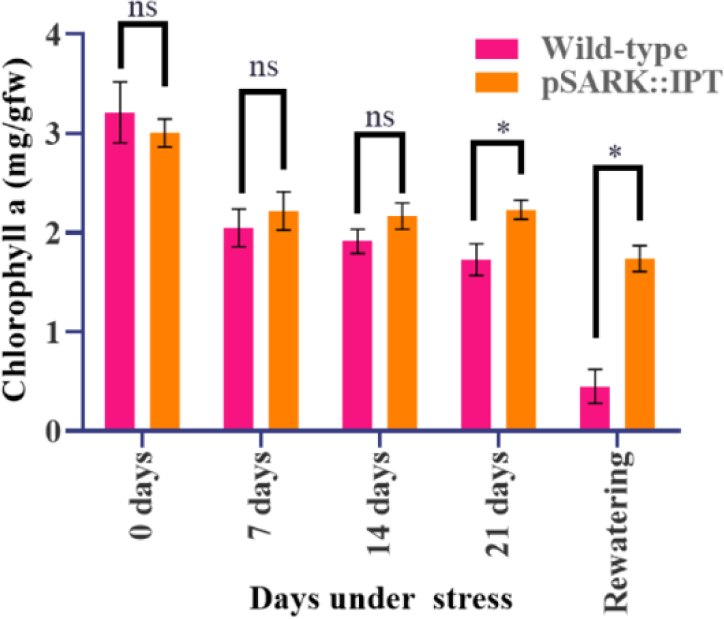
Comparison of chlorophyll a from leaves between wild type and pSARK::IPT plants at different time points. The results (n=15) are expressed as the mean ±SE. Bars/lines marked with asterisk (*) indicate that their means are statistically different from each other at 95% confidence interval.

Although chlorophyll b (the accessory pigment) was higher in pSARK::IPT maize, it did not differ significantly (p>0.05) from that of the wild type up to 14^th^ day of the stress period (Fig 6). However, significant differences (p<0.05) were observed after 21 days of stress (3.80mg/gfw in pSARK::IPT vis a vis 2.47mg/gfw in wild type plants). Moreover, chlorophyll b was significantly (p<0.05) three times higher in pSARK::IPT than in wild type plants after rewatering for three days. Regarding the ratio of chlorophyll a to chlorophyll b, there was no significant difference (p<0.05) between the two genotypes throughout the stress period.

**Fig 6:**
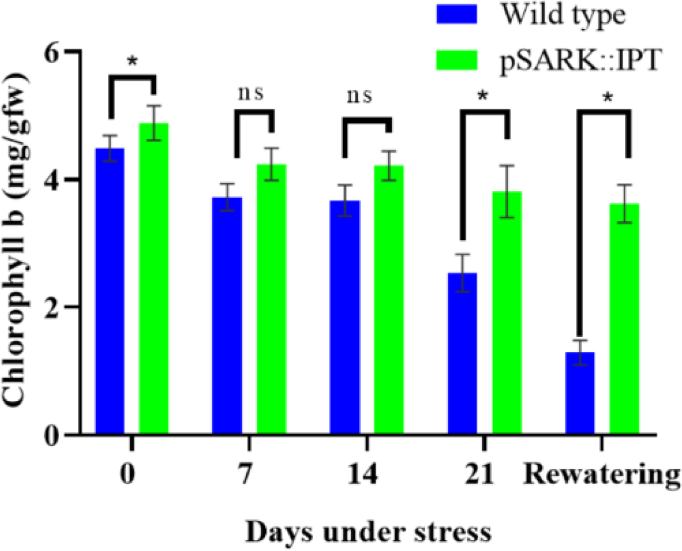
Comparison of chlorophyll b between pSARK::IPT and wild type plants. The results (n=15) are expressed as the mean ±SE. Bars/lines marked with asterisk (*) indicate that their means are statistically different from each other at 95% confidence interval.

Notably, the ratio of chlorophyll a to chlorophyll b increased after 21 days of stress in both groups from 0.4825 to 0.6509 in wild type and from 0.4977 to 0.6565 in pSARK::IPT (Fig 7).

**Fig 7:**
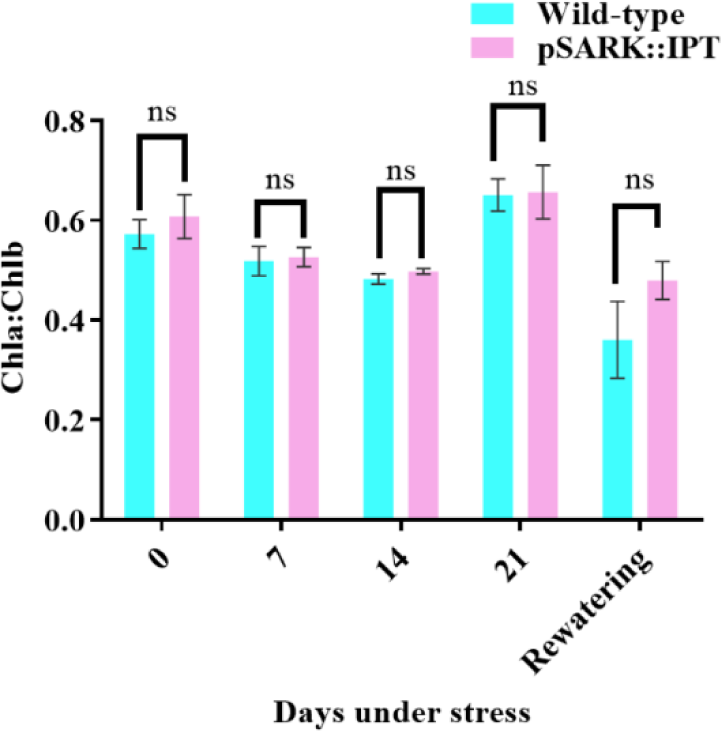
Chlorophyll a to chlorophyll b ratios comparison between pSARK::IPT and wild type plants under drought stress. The results (n=15) are expressed as the mean ±SE. Bars/lines marked with asterisk (*) indicate that their means are statistically different from each other confidence interval.

### Effect of drought on antioxidant enzymes activity

The CAT activity was three times higher in pSARK::IPT plants under drought stress than in well-watered conditions and this was significant. On the other hand, CAT activity did not differ significantly in wild type plants under both well-watered and stressed conditions. Moreover, CAT activity was 4 times higher in pSARK::IPT plants than in wild type plants under drought stress (Fig 8) and this was considered significant at p<0.05. The Ascorbate peroxidase (APX) activity was 6 and 3 times higher under drought stress than under well-watered conditions in pSARK::IPT and wild type plants respectively. Moreover, the enzyme activity was three times higher in pSARK::IPT than in wild type plants under drought stress (Fig 8).

**Fig 8:**
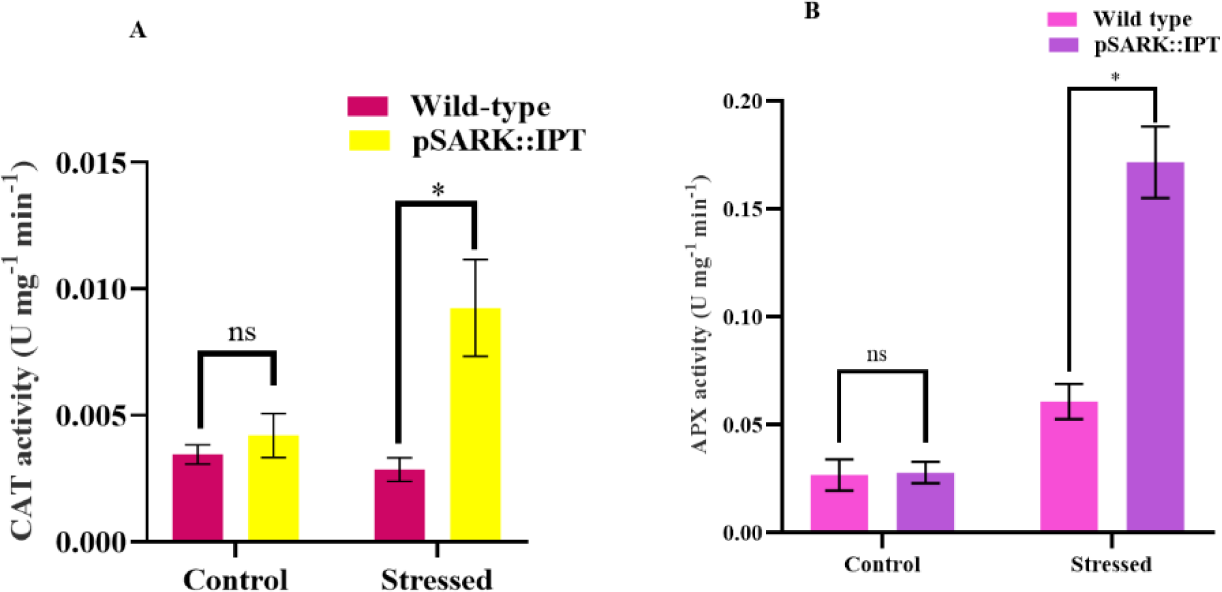
Antioxidant enzyme activities under well-watered and stressed conditions. **A**: CAT enzyme activity. **B**: APX enzyme activity. The results (n=10) are expressed as the mean ±SE. Asterisks (*) indicate a statistical significance at p<0.05.

### Effect of drought on plant height, leaf number and ASI

Plant height and leaf number per plant were recorded at the end of the stress period indicated that the average height of pSARK::IPT plants was significantly higher (p<0.05) than wild type plants under drought stress. Under drought stress, both wild type and pSARK::IPT plants had the same number of leaves per plant as shown in Table 1. Plants under drought stress tasseled two weeks after the well-watered plants. Regarding ASI, none of the wild type plants silked under drought stress. Most plants under drought stress produced anthers with little or no pollen. The ASI was 13 days on average in pSARK::IPT under drought stress which was significantly lower (p<0.05) than that of the wild type plants at 30 days (Table 1).

**Table 1.** Agronomic trait parameters under drought stress.

| Trait | Wild-type | pSARK::IPT | P-value |
| --- | --- | --- | --- |
| ASI | 13.40±1.628 | 29.89±1.033* | <0.0001 |
| Leaf number | 8.750±0.4163 | 8.800±0.4163 | 0.934 |
| Plant height | 70.25±3.085 | 87.70±1.880* | 0.0002 |

The results (n=10) are expressed as the mean ±SE. Asterisks (\*) indicate a statistical significance at p<0.05.

However, no significant variation (p>0.05) in height was observed between the two groups under well-watered conditions as shown in Table 2. On average, stressed plants were significantly (p<0.05) shorter than well-watered controls. while in well-watered conditions wild type plants had fewer leaves per plant (8) that did not differ significantly (p>0.05) from pSARK::IPT plants which had 10 leaves per plant (Table 1 and Table 2). On the other hand, ASI was five days in both wild type and pSARK::IPT conditions under well-watered conditions (Table 2).

**Table 2.** Agronomic trait parameters under well-watered conditions.

| Trait | Wild-type | pSARK::IPT | P-value |
| --- | --- | --- | --- |
| ASI | 4.600±0.3209 | 4.600±0.3625 | >0.9999 |
| Leaf number | 8.667 ± 0.3327 | 10.56 ± 0.3379* | 0.0017 |
| Plant height | 107.6 ± 7.627 | 112.9 ± 4.441 | 0.5338 |

The results (n=10) are expressed as the mean ±SE. Asterisks (\*) indicate a statistical significance at p<0.05

### Effect of drought on yield

Pollination was severely affected under drought stress. The tassels had little pollen. In most cases, pollen mortality was evident. The wild type plants had no ears since they didn’t silk while pSARK::IPT plants had ears with no kernels (Fig 9). Moreover, under well-watered conditions, the average seed weight (weight of 10 seeds) was significantly higher (p<0.05) in pSARK::IPT plants than in wild type plants (Table 3). Additionally, pSARK::IPT plants had significantly more seeds per comb than the wild type plants (Table 3)

**Fig 9.**
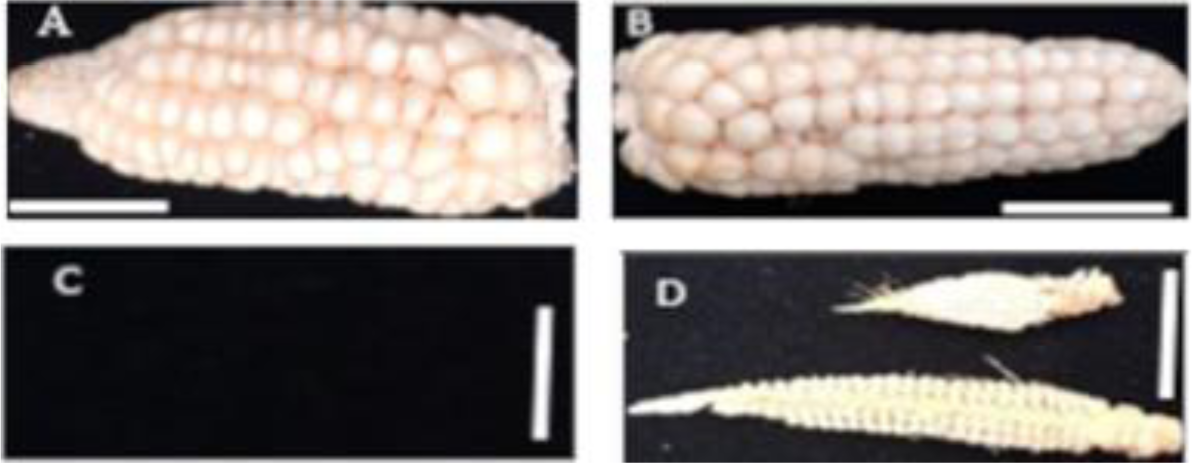
Comparison of yield and yield components between pSARK::IPT and wild type plants under drought stress and well-watered conditions. **A** and **B** Wild type and pSARK::IPT (**B**) ears under well-watered conditions. **C** and **D** represent wild type and pSARK::IPT ears under drought stress respectively. Scale bar = 1cm

**Table 3:**
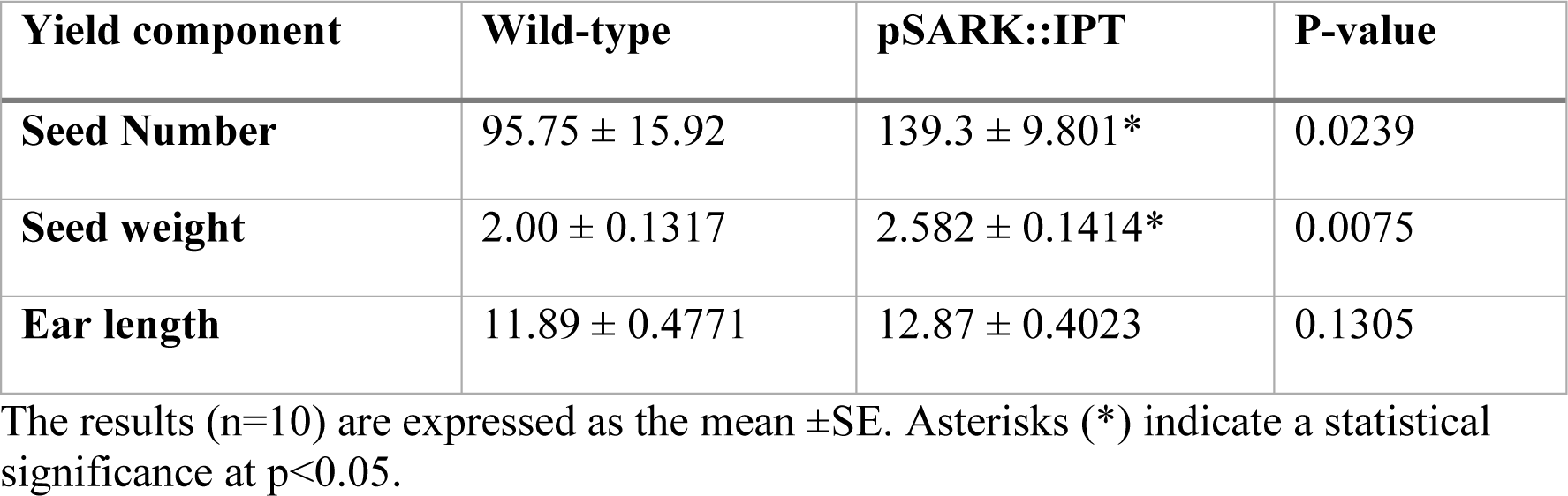
Comparison of yield and yield components between pSARK::IPT and wild type plants under well-watered conditions.

| Yield component | Wild-type | pSARK::IPT | P-value |
| --- | --- | --- | --- |
| Seed Number | 95.75 ± 15.92 | 139.3 ± 9.801* | 0.0239 |
| Seed weight | 2.00 ± 0.1317 | 2.582 ± 0.1414* | 0.0075 |
| Ear length | 11.89 ± 0.4771 | 12.87 ± 0.4023 | 0.1305 |

The results (n=10) are expressed as the mean ±SE. Asterisks (\*) indicate a statistical significance at p<0.05.

### Drought effect on shoots and roots

The primary roots were significantly longer (p<0.05) in pSARK::IPT than in wild type plants under drought stress. Moreover, the number of seminal roots was 1.4 times significantly higher (p<0.05) in pSARK::IPT than in wild type maize under drought stress as shown in Table 4. There was no significant difference (p<0.05) in shoot and root weights between pSARK::IPT and wild type plants under drought stress (Table 4 and Fig 10)

**Table 4:** Root parameters measured in wild type and pSARK::IPTunder drought stress.

| Root parameter | Wild-type | pSARK::IPT | P-value |
| --- | --- | --- | --- |
| Number of seminal roots | 22.00±1.711 | 29.60±1.586* | 0.0043 |
| Root length | 26.91±1.436 | 31.10 ±1.274* | 0.0352 |
| Biomass | 20.12±2.067 | 22.04±1.768 | 0.4882 |
| Fresh root weight | 9.965±1.638 | 9.582±1.349 | 0.8602 |
| Dry root weight | 2.7653±0.2944 | 3.393±0.2196 | 0.1045 |
| Fresh shoot weight | 47.14±6.860 | 62.15±6.867 | 0.1394 |
| Dry shoot weight | 16.79±2.899 | 18.58±1.849 | 0.5963 |

The results (n=10) are expressed as the mean ±SE. Asterisks (\*) indicate a statistical significance at p<0.05.

Under well-watered conditions, the number of seminal roots in pSARK::IPT plants was higher than that of the wild type plants. Additionally, the primary roots were longer in pSARK::IPT maize than in wild type maize but it was not statistically significant (p>0.05). Root and shoot fresh weights were significantly (p<0.05) 10 and 8 times higher under well-watered conditions than under drought stress respectively. Shoot fresh weight in pSARK::IPT was 1.3 times higher than wild type under well-watered conditions albeit non-significant as shown in Table 5.

**Table 5.** Root traits measured in wild type and pSARK::IPT under well-watered conditions.

| Root parameter | Wild-type | pSARK::IPT | P-value |
| --- | --- | --- | --- |
| Number of seminal roots | 30.22 ± 1.479 | 40.00 ± 2.442* | 0.0031 |
| Root length | 48.44 ± 2.450 | 50.89 ± 3.710 | 0.59 |
| Biomass | 147.04±23.10 | 167.8±12.26 | 0.446 |
| Fresh root weight | 107.5 ± 22.72 | 96.16 ± 17.39 | 0.8602 |
| Dry root weight | 34.56 ± 4.072 | 25.10 ± 5.241 | 0.1824 |
| Fresh shoot weight | 423.3 ± 26.80 | 429.1 ± 50.69 | 0.9208 |
| Dry shoot weight | 131.5 ± 11.84 | 122.3 ± 20.81 | 0.704 |

The results (n=10) are expressed as the mean ±SE. Asterisks (\*) indicate a statistical significance at p<0.05.

### Nutritional composition of transgenic maize

Upon harvesting the maize and quantifying yield, food component analysis of the mature kernels revealed that crude fat was significantly higher (p<0.05) in pSARK::IPT (13.82%) than in wild type (5.59%). Moreover, the crude protein content was significantly lower (p<0.05) in wild type (3.18%) than in pSARK::IPT (5.69%) as shown in Fig 11. Moisture content was higher in pSARK::IPT maize than in the wild type albeit non significantly (p>0.05). Furthermore, Crude fibre and ash content did not differ significantly between the wild type and pSARK::IPT.

**Fig 11:**
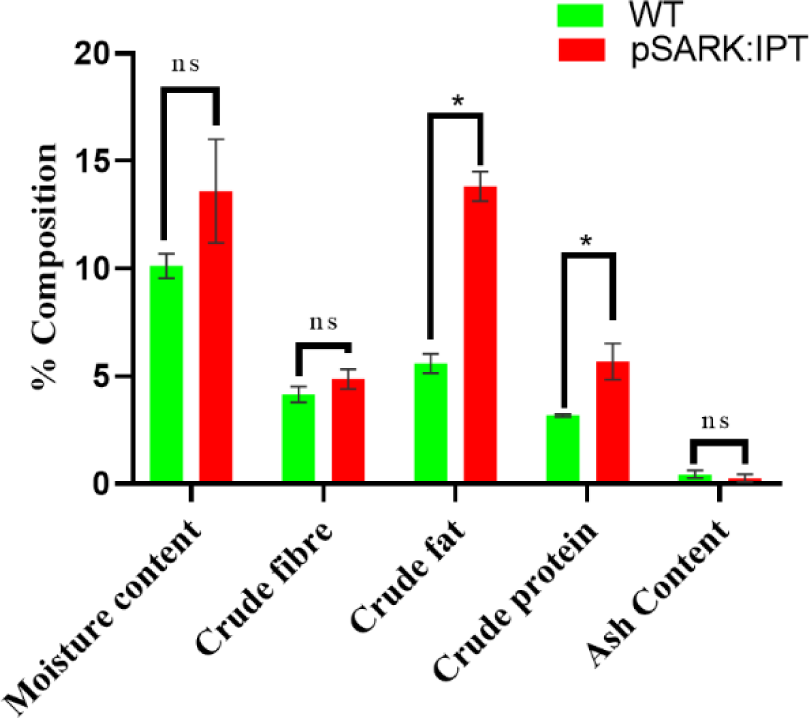
Proximate analysis of kernels of wild type and pSARK::IPT maize under well-watered conditions. The results (n=3) are expressed as the mean ±SE. Asterisks (*) indicate a statistical difference at p<0.05

## Discussion

One of the key challenges in developing successful transgenic crops is achieving stable and consistent expression of the inserted transgene in the plant’s cells across generations. Inconsistent expression can lead to unpredictable and unreliable results in terms of the desired trait expression. Evaluating the nutritional composition of transgenic plants is a crucial step prior to the approval of any transgenic event. Various nucleic acids and proteins play a pivotal role in governing gene expression. These biomolecules dictate whether a gene undergoes transcription and if this transcription results in the production of a protein that manifests as a specific phenotype.

The study established the persistence of the pSARK::IPT gene across multiple generations of maize plants (T3 and T4). This is a crucial aspect for transgenic crop development, as stable and consistent expression of the inserted gene ensures reliable trait expression over time. The findings suggest that the pSARK::IPT gene can maintain its presence and impact through subsequent generations, contributing to the sustainability of desired traits.

Furthermore, the transgenic maize showed improved drought tolerance depicted in its high relative water content, delayed leaf senescence, higher chlorophyll content and increased antioxidant enzyme activity when compared to the wild type controls. The balance between water supply to the leaf and its loss through transpiration and evaporation is well reflected by the relative water content (46). Chlorophyll degradation is a good indicator of senescence. The high chlorophyll content resulted from a delay in leaf senescence, a key function of cytokinins. Visible color change is a key characteristic of senescing leaves, which is closely linked to chlorophyll degradation and concomitant synthesis of anthocyanins and carotenoids (47).

Drought stress causes an imbalance in the production of reactive oxygen species (ROS) and its scavenging through the antioxidant defense system (48–50). Antioxidant enzymes are the most effective in the scavenging of ROS (51,52). Several studies have confirmed multiple correlations between antioxidant enzyme activities and CKs (18,53,54). CKs may act transcriptionally to alter responses to ROS often produced during plant abiotic stress (55). Glutathione peroxidase (GPX), APX and CAT are the primary ROS scavengers in plants (56)

This study was designed in such a way that the stress period coincided with the pre-anthesis stage and persisted for 21 days, which is considered a long-term drought (31). We found that the water deficit treatment affected flowering, grain yield and kernel set of the pSARK::IPT and wild type maize plants. Flowering was delayed and embryo aborted in pSARK::IPT maize, whereas the wild type maize did not produce silk at all. Anthesis-silking interval was longer in stressed plants compared to well-watered ones.

Drought stress severely affects maize yield when it occurs at the flowering stage (57–59). Gametogenesis, fertilization, and embryogenesis are severely affected by drought, limiting seed development thus lowering crop yields (60). (61) reported pollen sterility as a common symptom during drought stress. It reduces pollen germination as well as inhibits growth of the pollen tube consequently impairing fertilization (62,63). Silk is more susceptible to drought stress than tassel causing delay in silk emergence (64).

Drought inhibits photosynthesis by reducing the capacity for Ribulose-1,5-bisphosphate regeneration as well as its carboxylation efficiency. Whole-plant senescence severely affects seed filling (65). Water deficit in the course of seed development decreases the number of amyloplasts and endosperm cells, consequently reducing kernel/seed sink strength (66). This negatively impacts the rate and duration at which endosperm gathers starch which ultimately reduces grain weight (67). Ovary abortion occurs due to the increase in non-reducing sugars as well as the inability to accumulate starch under water deficit which consequently decreases grain yield (68,69). Moreover, drought during seed filling causes early senescence and reduces seed-filling duration and enhances assimilate remobilization from the source to sink (64).

Cytokinins (CKs) stimulate the endosperm cells to divide rapidly, which enhances grain filling (70). Increased CKs levels may promote sink strength via up-regulating genes involved in cell division. This can be through sugar signaling, that involves increased phloem unloading and sugar import to endospermic cells via a cell wall-associated enzyme invertase (64). The study highlights the role of cytokinins in stimulating endosperm cell division and enhancing grain filling. This knowledge can be leveraged for optimizing seed development and grain yield. By understanding the impact of pSARK::IPT on endosperm growth, researchers and breeders can target strategies to improve the efficiency of nutrient accumulation and seed filling.

Roots are the first plant organs to be exposed to the drying soil hence the primary sensors/detectors of drought stress (71). The structural root traits affected by drought include volume, density, length and number and these ultimately limit the functioning of the entire plant (71,72). The observed increase in seminal roots and higher shoot fresh weight in transgenic maize plants under well-watered conditions suggests that the pSARK::IPT gene might influence root architecture and growth dynamics. This has implications for overall plant health and resource acquisition, potentially contributing to enhanced performance and productivity.

Interestingly, pSARK::IPT plants performed better than wild type under well-watered conditions. More seminal roots were recorded in pSARK::IPT maize than in wild type maize plants. Furthermore, the fresh weight of shoots in pSARK::IPT plants was non-significantly higher compared to that of wild type under well-watered conditions. No significant variations were recorded on dry weight between the studied genotypes. This could be due to water evaporation from the plant during drying reducing variations observed in fresh weights. It has been reported that maize transformed by particle bombardment with pSARK::IPT construct had similar biomass under both stressed and well-watered conditions (30).

The current study further established that the pSARK::IPT plants showed significantly higher seed number and seed weight than their wild type counterparts under well-watered conditions. The observation that pSARK::IPT plants performed better than wild-type plants under well-watered conditions is intriguing. This suggests that the genetic modification may have conferred some advantage in terms of water utilization, nutrient uptake, or growth processes. Further investigation into the specific mechanisms behind this improved performance would be valuable. The pSARK::IPT gene has not been reported to improve the yield of maize under well-watered conditions. However, (73) reported improved yields in canola plants when *IPT* was driven by a developmentally-induced promoter AtMYB32xs.

This study reported that under well-watered conditions, crude fat was significantly higher in pSARK::IPT maize than in wild type maize. Likewise, crude protein content was significantly lower in wild type maize than in pSARK::IPT maize (Fig 11). Moisture content was higher in pSARK::IPT maize than in the wild type albeit non-significantly possibly due to increased seed filling duration in pSARK::IPT maize. The longer duration might allow for more water uptake, leading to slightly higher moisture content in the transgenic maize. The altered nutritional composition of transgenic maize, including higher crude fat content and possibly moisture content, has potential implications for both animal feed and human consumption. These changes may impact the nutritional quality and utility of the maize crop in various agricultural and food-processing contexts.

Conversely, it was reported that oil from peanuts transformed with pSARK::IPT had the same quality as that of wild type under an environment that was well-watered(74). However, transgenic (pSARK::IPT) rice plants had a 15 % increase in grain starch when they were maintained in a well-watered environment (75).

Studies have reported that genes that code for enzymes of the starch degradation pathway were downregulated in pSARK::IPT transformed rice while those that encode protein kinases Osk24(Os08g37800) and Osk1 (Os05g45420) were upregulated when plants were grown in a well-watered environment (75). The role of Osk1 is to unload sucrose through the sucrose synthase pathway while Osk24 plays a fundamental role in the conversion of sucrose to starch (76).

Sugars are the respiratory substrates for the production of energy and metabolites required to synthesize macromolecules. Transcriptome analysis of sucrose-deprived rice suspension cells reported a decline in the expression of transcripts associated with synthesis of fatty acid vis-a vis an increase in those involved in degradation of fatty acid such as fatty acid multifunctional proteins, 3-ketoacyl-CoA thiolase and acyl-CoA oxidase (77). The ultimate size and number of the endosperm cells determines the amount of proteins and starch that accumulates in the grains which are affected by the duration and rate of the grain filling process (75). Cytokinins are associated with strengthening of the sink/source relationship in pSARK::IPT plants which consequently leads to higher yields (78).

The study’s evaluation of the pSARK::IPT gene’s effects spans a limited time frame (21 days of drought stress) and extreme drought conditions. It does not fully explore how the gene interacts with varying levels of stress severity. The study describes observed effects of cytokinins on various traits but does not delve deeply into the underlying molecular mechanisms driving these changes. A more comprehensive understanding of the genetic and biochemical pathways involved could provide more insights.

## Conclusion

This study presumed that the SARK promoter is not only induced by drought but also by maturation which elevates the levels of cytokinin under well-watered conditions. The study further hypothesize that the better performance of pSARK::IPT under well-watered conditions could be due to downregulation of enzymes encoding the starch degradation pathway as well as upregulation of SnRK1-type of protein kinase genes. Besides, the activation of cell wall invertase enzyme that enhances translocation of plant nutrients from the source to the sink and increased cell division in the endosperm, which are associated with increased cytokinin, could result in enhanced performance in the transgenic maize.

The study demonstrates that transgenic maize expressing the pSARK::IPT gene exhibited improved drought tolerance compared to wild-type plants. This finding suggests that manipulating cytokinin levels through gene expression can lead to better water retention, delayed senescence, and higher antioxidant enzyme activity. These traits collectively contribute to the plant’s ability to withstand prolonged water deficits, which is a significant advantage for agricultural productivity in regions prone to drought.

While the study contributes valuable information about the potential benefits of the pSARK::IPT gene in transgenic maize, it is essential to recognize its limitations. Further research, including field trials, in-depth molecular analyses, and assessments across diverse genotypes and environmental conditions, is necessary to fully understand the broader implications and potential challenges associated with using this gene for crop improvement.

## Acknowledgments

The authors would like to express their gratitude to Dr. Leta Tulu Bedada and his team for their preliminary research, which laid the foundation for this study.

## Author Contributions

Conceptualization: Richard Oduor, Mathew Ngugi

Funding acquisition: Richard Oduor, Mathew Ngugi

Investigation: Rose Mweru.

Resources: Richard Oduor, Mathew Ngugi

Supervision: Richard Oduor, Mathew Ngugi

Writing – original draft: Rose Mweru, Shem B. Nchore

## References

1. Doebley J. The genetics of maize evolution. Vol. 38, Annual Review of Genetics. Annual Reviews; 2004. p. 37–59.

2. Byakod M. Evaluation of maize genotypes for moisture stress condition. 2017.

3. Sharma K, Misra RS. Molecular approaches towards analyzing the viruses infecting maize (Zea mays L.). J Gen Mol Virol. 2011;3(1):1–17.

4. Ranum P, Peña-Rosas JP, Garcia-Casal MN. Global maize production, utilization, and consumption. Ann N Y Acad Sci. 2014;1312(1):105–12.

5. Pardey PG, Andrade RS, Hurley TM, Rao X, Liebenberg FG. Returns to food and agricultural R&D investments in Sub-Saharan Africa, 1975–2014. Food Policy. 2016 Dec 1;65:1–8.

6. Awata LA, Tongoona P, Danquah E, Ifie BE, Suresh LM, Jumbo MB, et al. Understanding tropical maize (Zea mays L.): The major monocot in modernization and sustainability of agriculture in sub-Saharan Africa. Ijaar. 2019;7:32–77.

7. Daly J, Hamrick D, Gereffi G, Guinn A. Maize value chains in East Africa.

8. Him-Gonzalez C. Water resources for agriculture and food production.

9. Vörösmarty CJ, Green P, Salisbury J, Lammers RB. Global water resources: Vulnerability from climate change and population growth. Science (1979). 2000 Jul 14;289(5477):284–8.

10. Roser M. Two centuries of rapid global population growth will come to an end. Our World in Data. 2019 Jun 18;

11. Maltsoglou, Irini and Khwaja Y. Environment and natural resources management working paper. The BEFS Analysis for Tanzania Bioenergy. 2010;248.

12. Birch I. Economic growth in the arid and semi-arid lands of Kenya Question What recommendations have been made by reputable experts to support long-term sustainable economic growth in the Arid and Semi-Arid Lands of Kenya? 2018.

13. Kar RK. Plant responses to water stress: Role of reactive oxygen species. Plant Signal Behav. 2011 Nov;6(11):1741–5.

14. Anjum SA, Wang LC, Farooq M, Hussain M, Xue LL, Zou CM. Brassinolide application improves the drought tolerance in maize through modulation of enzymatic antioxidants and leaf gas exchange. J Agron Crop Sci. 2011 Jun;197(3):177–85.

15. Basu S, Ramegowda V, Kumar A, Pereira A. Plant adaptation to drought stress [version 1; referees: 3 approved]. F1000Res. 2016;5(July).

16. Akiyoshi DE, Klee H, Amasino RM, Nester EW, Gordon MP. T-DNA of Agrobacterium tumefaciens encodes an enzyme of cytokinin biosynthesis. Proc Natl Acad Sci U S A. 1984;81(19 I):5994–8.

17. Barry GF, Rogers SG, Fraley RT, Brand L. Identification of a cloned cytokinin biosynthetic gene. Proceedings of the National Academy of Sciences. 1984 Aug 181(15):4776–80.

18. Hai NN, Chuong NN, Tu NHC, Kisiala A, Hoang XLT, Thao NP. Role and regulation of cytokinins in plant response to drought stress. Plants. 2020;9(4):10–2.

19. Guo Y, Gan SS. Translational research on leaf senescence for enhancing plant productivity and quality. J Exp Bot. 2014;65(14):3901–13.

20. Sade N, Del Mar Rubio-Wilhelmi M, Umnajkitikorn K, Blumwald E. Stress-induced senescence and plant tolerance to abiotic stress. Jomo Kenyatta University of Agriculture and Technology user on. 2018;69(4):845–53.

21. Gregersen PL, Culetic A, Boschian L, Krupinska K. Plant senescence and crop productivity. Vol. 82, Plant Molecular Biology. 2013. p. 603–22.

22. Joshi S, Choukimath A, Isenegger D, Panozzo J, Spangenberg G, Kant S. Improved Wheat Growth and Yield by Delayed Leaf Senescence Using Developmentally Regulated Expression of a Cytokinin Biosynthesis Gene. Front Plant Sci. 2019 Oct 18;10(October):1–11.

23. Ahuja MR, Fladung M. Integration and inheritance of transgenes in crop plants and trees. Vol. 10, Tree Genetics and Genomes. Springer Verlag; 2014. p. 779–90.

24. Bandopadhyay R, Haque I, Singh D, Mukhopadhyay K. Levels and stability of expression of transgenes. In: Transgenic Crop Plants. Springer-Verlag Berlin Heidelberg; 2010. p. 145–86.

25. Yi DX, Fang ZY, Yang LM. Inheritance and expression of Bt cry1Ba3 gene in progeny from transformed cabbage plants. Mol Biol Rep. 2020 Apr 1;47(4):2583–9.

26. Erasmus R, Pieters R, Plessis H Du, Hilbeck A, Trtikova M, Erasmus A, et al. Introgression of a cry1Ab transgene into open pollinated maize and its effect on Cry protein concentration and target pest survival. PLoS One. 2019 Dec 1;14(12).

27. LIU H yun, Wang K, Wang J, Du L Pu, Pei X Wu, Ye X Guo. Genetic and agronomic traits stability of marker-free transgenic wheat plants generated from Agrobacterium-mediated co-transformation in T2 and T3 generations. J Integr Agric. 2020 Jan 1;19(1):23–32.

28. Zhao Y, Guo L, Wang H, Huang D. Integration and expression stability of transgenes in hybriding transmission of transgenic rice plants produced by particle bombardment. Mol Plant Breed. 2011.

29. Robson PRH, Donnison IS, Wang K, Frame B, Pegg SE, Thomas A, et al. Leaf senescence is delayed in maize expressing the Agrobacterium IPT gene under the control of a novel maize senescence-enhanced promoter. Plant Biotechnol J. 2004;2(2):101–12.

30. Décima Oneto C, Otegui ME, Baroli I, Beznec A, Faccio P, Bossio E, et al. Water deficit stress tolerance in maize conferred by expression of an isopentenyltransferase (IPT) gene driven by a stress- and maturation-induced promoter. J Biotechnol. 2016 Feb 20;220:66–77.

31. Leta TB, Miccah SS, Steven MR, Wondyifraw T, Charless M, Clet WM, et al. Drought tolerant tropical maize (Zea mays L.) developed through genetic transformation with isopentenyltransferase gene. Afr J Biotechnol. 2016 15(43):2447–64.

32. Tamari F, Hinkley CS, Ramprashad N. A Comparison of DNA extraction methods using Petunia hybrida Tissues. 2013.

33. Lorenz TC. Polymerase chain reaction: Basic protocol plus troubleshooting and optimization strategies. Journal of Visualized Experiments. 2012 May 22;63(63):3998.

34. Rathod SD, Dahiwalkar SD, Gorantiwar SD, Kamble BM, Shinde MG. Physical properties of waterlogged vertisols under subsurface drainage system with different drain spacings and depths. International Journal of Agricultural Engineering. 2017;10(1):22–30.

35. Miccah SS, Leta TB, Steven MR, Mathew PN, Revel I, Jennifer AT, et al. Drought tolerance in transgenic tropical maize (Zea mays L.) by heterologous expression of peroxiredoxin2 gene-XvPrx2. Afr J Biotechnol. 2016;15(25):1350–62.

36. Soltys-Kalina D, Plich J, Strzelczyk-Żyta D, Śliwka J, Marczewski W. The effect of drought stress on the leaf relative water content and tuber yield of a half-sib family of ‘Katahdin’-derived potato cultivars. Breed Sci. 2016 Apr 8;66(2):328–31.

37. Kumari R, Ashraf S, Khatik SK, Bagdi DL, Bagri GK, Bagri DK. Extraction and estimation of chlorophyll content of seed treated lentil crop using DMSO and acetone. ∼ 249 ∼ Journal of Pharmacognosy and Phytochemistry. 2018;7(3):249–50.

38. Wellburn AR. The spectral determination of chlorophylls a and b, as well as total carotenoids, using various solvents with spectrophotometers of different resolution. J Plant Physiol. 1994;144(3):307–13.

39. Elavarthi S, Martin B. Spectrophotometric assays for antioxidant enzymes in plants. Methods Mol Biol. 2010;639:273–81.

40. Senthilkumar M, Amaresan N, Sankaranarayanan A. Estimation of Ascorbate Peroxidase (APX). In Humana, New York, NY; 2021. p. 119–21.

41. Bradley RL. Moisture and Total Solids Analysis. In 2010. p. 85–104.

42. Marshall MR. Ash Analysis. In 2010. p. 105–15.

43. Min DB, Ellefson WC. Fat Analysis. In 2010. p. 117–32.

44. Möller J. Comparing methods for fibre determination in food and feed. Foss. 2014;(1):1–6.

45. Nielsen SS. Food Analysis Laboratory Manual. Cham: Springer International Publishing; 2017. (Food Science Text Series).

46. Lugojan C, Ciulca S. Analysis of excised leaves water loss in winter wheat. Vol. 15, Forestry and Biotechnology. 2011.

47. Zhao MH, Li X, Zhang XX, Zhang H, Zhao XY. Mutation mechanism of leaf color in plants: A review. Vol. 11, Forests. MDPI AG; 2020.

48. Choudhury FK, Rivero RM, Blumwald E, Mittler R. Reactive oxygen species, abiotic stress and stress combination. Plant Journal. 2017 Jun 1;90(5):856–67.

49. Verma G, Srivastava D, Tiwari P, Chakrabarty D. ROS modulation in crop plants under drought stress. In: Reactive Oxygen, Nitrogen and Sulfur Species in Plants: Production, Metabolism, Signaling and Defense Mechanisms. Taylor and Francis; 2019. p. 311–36.

50. Wei T, Wang Y, Xie Z, Guo D, Chen C, Fan Q, et al. Enhanced ROS scavenging and sugar accumulation contribute to drought tolerance of naturally occurring autotetraploids in Poncirus trifoliata. Plant Biotechnol J. 2019 Jul 1;17(7):1394–407.

51. Saruhan N, Saglam A, Kadioglu A. Salicylic acid pretreatment induces drought tolerance and delays leaf rolling by inducing antioxidant systems in maize genotypes. Acta Physiol Plant. 2012 Jan;34(1):97–106.

52. Osmolovskaya N, Shumilina J, Kim A, Didio A, Grishina T, Bilova T, et al. Methodology of drought stress research: Experimental setup and physiological characterization. Int J Mol Sci. 2018;19(12).

53. de Moura FB, Marcos MR, do N. Simões A, da Silva SLF, de Medeiros DC, de A. Paes R, et al. Participation of cytokinin on gas exchange and antioxidant enzymes activities. Vol. 22, Indian Journal of Plant Physiology. Springer Verlag; 2017. p. 16–29.

54. Wu X, Zhu Z, Li X, Zha D. Effects of cytokinin on photosynthetic gas exchange, chlorophyll fluorescence parameters and antioxidative system in seedlings of eggplant (Solanum melongena L.) under salinity stress. Acta Physiol Plant. 2012 Nov;34(6):2105–14.

55. Zwack PJ, Rashotte AM. Cytokinin inhibition of leaf senescence. Plant Signal Behav. 2013 Jul;8(7):1–7.

56. Laxa M, Liebthal M, Telman W, Chibani K, Dietz KJJ. The role of the plant antioxidant system in drought tolerance. Antioxidants. 2019;8(4).

57. Meseka S, Menkir A, Bossey B, Mengesha W. Performance assessment of drought tolerant maize hybrids under combined drought and heat stress. Agronomy. 2018 Nov 22;8(12):274.

58. Jain M, Kataria S, Hirve M, Prajapati R. Water Deficit Stress Effects and Responses in Maize. 2019;129–51.

59. Sah RP, Chakraborty M, Prasad K, Pandit M, Tudu VK, Chakravarty MK, et al. Impact of water deficit stress in maize: Phenology and yield components. Sci Rep. 2020 Dec 1;10(1):1–15.

60. Farooq M, Hussain M, Siddique KHM. Drought Stress in Wheat during Flowering and Grain-filling Periods. CRC Crit Rev Plant Sci. 2014;33(4):331–49.

61. Dong B, Zheng X, Liu H, Able JA, Yang H, Zhao H, et al. Effects of drought stress on pollen sterility, grain yield, abscisic acid and protective enzymes in two winter wheat cultivars. Front Plant Sci. 2017 Jun 20;8.

62. Fang X, Turner NC, Yan G, Li F, Siddique KHM. Flower numbers, pod production, pollen viability, and pistil function are reduced and flower and pod abortion increased in chickpea (Cicer arietinum L.) under terminal drought. J Exp Bot. 2010 Jan 1;61(2):335–45.

63. Gusmao M, Siddique KHM, Flower K, Nesbitt H, Veneklaas EJ. Water Deficit during the Reproductive Period of Grass Pea (Lathyrus sativus L.) Reduced Grain Yield but Maintained Seed Size. J Agron Crop Sci. 2012 Dec;198(6):430–41.

64. Sehgal A, Sita K, Siddique KHM, Kumar R, Bhogireddy S, Varshney RK, et al. Drought or/and heat-stress effects on seed filling in food crops: Impacts on functional biochemistry, seed yields, and nutritional quality. Front Plant Sci. 2018;871(November):1–19.

65. Basuki K. 済無No Title No Title. ISSN 2502-3632 (Online) ISSN 2356-0304 (Paper) Jurnal Online Internasional & Nasional Vol 7 No1, Januari – Juni 2019 Universitas 17 Agustus 1945 Jakarta. 2019;53(9):1689–99.

66. Ndlovu E, van Staden J, Maphosa M. Morpho-physiological effects of moisture, heat and combined stresses on Sorghum bicolor [Moench (L.)] and its acclimation mechanisms. Plant Stress. 2021 Dec;2:100018.

67. Fábián A, Jäger K, Rakszegi M, Barnabás B. Embryo and endosperm development in wheat (Triticum aestivum L.) kernels subjected to drought stress. Plant Cell Rep. 2011 Apr;30(4):551–63.

68. Jiang Z, Piao L, Guo D, Zhu H, Wang S, Zhu H, et al. Regulation of maize kernel carbohydrate metabolism by abscisic acid applied at the grain-filling stage at low soil water potential. Sustainability (Switzerland). 2021 Mar 2;13(6).

69. Khodaeiaminjan M, Bergougnoux V. Barley Grain Development during Drought Stress: Current Status and Perspectives. Cereal Grains. 2021 Apr 24;

70. Kong L, Guo H, Sun M. Signal transduction during wheat grain development. Planta. 2015;241(4):789–801.

71. Aslam M, Maqpool M, Cengiz R. Drought stress in maize (Zea Mays L.): effects, resistance mechanisms, global achievements, and biological strategies. Vol. 8, Springer Briefs in Agriculture. 2015. 5–18 p.

72. Comas L, Becker S, Cruz VM V., Byrne PF, Dierig DA. Root traits contributing to plant productivity under drought. Front Plant Sci. 2013 Nov 5;4(NOV):442.

73. Kant S, Burch D, Badenhorst P, Palanisamy R, Mason J, Spangenberg G. Regulated Expression of a cytokinin biosynthesis gene IPT delays leaf senescence and improves yield under rainfed and irrigated conditions in canola (Brassica napus L.). PLoS One. 2015 Jan 20;10(1).

74. Gregersen PL, Culetic A, Boschian L, Krupinska K. Plant senescence and crop productivity. Plant Mol Biol [Internet]. 2013 Aug 25 [cited 2023 Aug 14];82(6):603–22. Available from: https://link.springer.com/article/10.1007/s11103-013-0013-8

75. Peleg Z, Reguera M, Tumimbang E, Walia H, Blumwald E. Cytokinin-mediated source/sink modifications improve drought tolerance and increase grain yield in rice under water-stress. Plant Biotechnol J. 2011;9(7):747–58.

76. Raffan S, Oddy J, Halford NG. The sulphur response in wheat grain and its implications for acrylamide formation and food safety. Int J Mol Sci. 2020;21(11):1–20.

77. Wang HJ, Wan AR, Hsu CM, Lee KW, Yu SM, Jauh GY. Transcriptomic adaptations in rice suspension cells under sucrose starvation. Plant Mol Biol. 2007 Mar;63(4):441– 63.

78. Wang W, Hao Q, Tian F, Li Q, Wang W. Cytokinin-Regulated Sucrose Metabolism in Stay-Green Wheat Phenotype. PLoS One. 2016;11(8).

